# The GspCD-dependent type II secretion system drives necrotizing soft tissue infection by *Aeromonas hydrophila*

**DOI:** 10.64898/2026.04.30.721992

**Authors:** Yuka Tonosaki, Kohei Yamazaki, Shota Owada, Kei Yamaguchi, Takashige Kashimoto

**Affiliations:** Laboratory of Veterinary Public Health, School of Veterinary Medicine, Kitasato University, Aomori, Japan

## Abstract

Necrotizing soft tissue infections (NSTIs) are fulminant bacterial diseases characterized by rapid tissue destruction, systemic deterioration, and high mortality. *Aeromonas hydrophila* is an important causative agent of NSTIs, but the system-level bacterial mechanisms that coordinate tissue destruction, in vivo expansion, dissemination, and host lethality remain incompletely understood. Here, we investigated the contribution of the GspCD-dependent type II secretion system (T2SS) to *A. hydrophila* pathogenesis using transposon mutants, extracellular protein analyses, and a mouse NSTI model. Mutants carrying transposon insertions in *gspD* and *gspC* showed defective secretion of a FLAG-tagged truncated AerA construct and markedly reduced hemolytic activity in culture supernatants. Comparative analysis of extracellular proteins further showed that disruption of *gspC* altered the extracellular protein landscape, with reduced abundance of multiple known or predicted virulence-associated factors, including AerA, Ahh, lipase, and metalloprotease. In the mouse NSTI model, both mutants exhibited attenuated virulence, including reduced serum markers of tissue injury, less severe histopathological damage, impaired in vivo expansion and dissemination, and decreased lethality. These defects were more pronounced in the *gspC* mutant than in the *gspD* mutant. Together, these findings show that the GspCD-dependent T2SS functions as a coordinated extracellular secretion system that drives tissue destruction, in vivo expansion, dissemination, and lethal outcome during *A. hydrophila* NSTI.

**IMPORTANCE:** Necrotizing soft tissue infections (NSTIs) are rapidly progressive, life-threatening bacterial infections, and *Aeromonas hydrophila* is an important causative agent. Here, we show that the GspCD-dependent type II secretion system (T2SS) drives *A. hydrophila* virulence in a murine NSTI model. Transposon mutants in *gspC* or *gspD* exhibited impaired extracellular protein secretion, reduced hemolytic activity, attenuated tissue damage, decreased bacterial proliferation and dissemination, and markedly reduced lethality. Comparative analysis further indicated that T2SS disruption alters the extracellular virulence landscape rather than affecting a single toxin alone. These findings provide in vivo evidence that coordinated T2SS-dependent secretion is a central determinant of severe *A. hydrophila* soft tissue infection.

## INTRODUCTION

Necrotizing soft tissue infections (NSTIs) are rapidly progressive and life-threatening bacterial diseases characterized by extensive destruction of subcutaneous tissue and skeletal muscle, systemic deterioration, and high mortality [1, 2]. *Aeromonas hydrophila* is an important etiologic agent of NSTIs, particularly in wound-associated infections following exposure to aquatic environments [3–7]. Clinical cases of *A. hydrophila* NSTIs are often fulminant and can rapidly progress to septic shock and death, underscoring the need to better understand the bacterial mechanisms that drive invasive soft tissue disease [4, 8, 9].

Several virulence factors have been implicated in the pathogenesis of *A. hydrophila*, including proteases, lipases, and extracellular toxins such as Aerolysin and hemolysin, many of which are secreted via the type II secretion system (T2SS) [5, 6, 10, 11]. Among these, Aerolysin has been extensively studied as a pore-forming toxin that damages host cell membranes and contributes to cytotoxicity [12–15]. However, the bacterial mechanisms that coordinate tissue destruction, bacterial proliferation within infected tissues, dissemination, and lethality during NSTIs remain incompletely understood. We have previously shown that motility is required for efficient spread of *A. hydrophila* within soft tissues [16], indicating that active bacterial dissemination is a key determinant of disease progression. Because this process is likely constrained by physical barriers such as the extracellular matrix and host cell integrity, we hypothesized that T2SS-dependent secretion of extracellular toxins and degradative enzymes promotes bacterial proliferation and dissemination by disrupting host tissues and removing these barriers. Thus, in addition to motility, secretion-dependent remodeling of the local tissue environment may be required for destructive and lethal NSTI progression.

The T2SS is a conserved secretion apparatus in Gram-negative bacteria that exports folded proteins from the periplasm to the extracellular milieu. T2SS substrates are first transported across the inner membrane by the Sec or Tat pathway and are then secreted across the outer membrane through a multiprotein complex containing the secretin GspD [17–20]. GspC is thought to contribute to substrate recognition and to the functional linkage of the secretion machinery [17–20]. In several Gram-negative pathogens, the T2SS is required for secretion of major virulence factors and hydrolytic enzymes, supporting a central role for this system in extracellular protein export and pathogenesis [17, 21–23]. Despite the established importance of secreted toxins in *Aeromonas* pathogenesis, the contribution of the T2SS itself to necrotizing soft tissue infection has not been directly defined in a murine model.

In the present study, we performed phenotypic screening of a transposon mutant library of *A. hydrophila* and identified mutants carrying insertions in *gspD* and *gspC*. Using these mutants, we investigated how the GspCD-dependent T2SS contributes to extracellular protein secretion, hemolytic activity, tissue damage, in vivo expansion and dissemination, and host lethality in a mouse NSTI model. Our findings indicate that this secretion system drives severe soft tissue infection by coordinating the export of extracellular factors that promote tissue destruction, bacterial spread, and host death.

## MATERIALS AND METHODS

### Bacterial strains and culture conditions

*Aeromonas hydrophila* clinical isolate RIMD 111065 and its derivatives were used in this study. The strain was originally isolated from human blood and was obtained from the Pathogenic Microbial Resource Unit, Research Institute for Microbial Diseases, Osaka University. A spontaneous rifampicin-resistant derivative of the wild-type (WT) strain was used for transposon mutagenesis and animal infection experiments. Bacteria were routinely cultured in Luria-Bertani (LB) broth at 37°C with shaking at 200 rpm or on LB agar plates. When required, chloramphenicol (5 µg/mL for *A. hydrophila* and 10 µg/mL for *Escherichia coli*) or kanamycin was added for plasmid maintenance or mutant selection.

For in vivo bioluminescence imaging, a bioluminescent derivative carrying the *luxCDABE* operon integrated into the chromosome was used. Chromosomal integration was achieved using a suicide vector pYAK1 [16], carrying the *luxCDABE* cassette flanked by *flaA* homologous arms, followed by allelic exchange. This chromosomal integration enabled stable maintenance of the bioluminescent marker without antibiotic selection.

### Plasmid construction and complementation

Genomic DNA from the WT strain was purified using the DNeasy Blood & Tissue Kit (Qiagen, Hilden, Germany). The *aerA*, *gspD*, and *gspC* genes were amplified by PCR using gene-specific primers.

For detection of Aerolysin secretion, a truncated *aerA* construct encoding the secretion signal and N-terminal region of AerA (104 amino acids) was amplified. A FLAG tag (8 amino acids) was fused to its C terminus, generating a construct with a predicted molecular mass of approximately 20 kDa. A truncated construct was used to avoid potential biological effects associated with expression of full-length Aerolysin.

The pACYC184 vector backbone was amplified by PCR and assembled with each insert using the NEBuilder HiFi DNA Assembly Master Mix (New England Biolabs, Ipswich, MA, USA). The resulting plasmids were transformed into *Escherichia coli* DH5α, and transformants were selected on LB agar plates containing chloramphenicol. Plasmids were then introduced into *A. hydrophila* mutant strains by conjugation, and complemented strains were selected on LB agar plates containing chloramphenicol.

### Construction of a transposon mutant library

A transposon mutant library was generated as described previously with minor modifications [24]. Briefly, *A. hydrophila* was statically cultured in LB broth at 25°C for more than 12 h, harvested by centrifugation at 10,000 × *g* for 1 min, and washed three times with lactose broth. *E. coli* BW19795 carrying pUT mini-Tn5 Km2 Tag was statically cultured in LB broth containing kanamycin at 37°C and washed in the same manner. The transposon was introduced into *A. hydrophila* by conjugation with BW19795, and transconjugants were selected on agar plates containing appropriate antibiotics.

### Identification of transposon insertion sites

Transposon insertion sites were identified by arbitrarily primed PCR as described previously [24]. Briefly, genomic DNA was extracted from each transposon mutant and subjected to PCR using a transposon-specific primer together with arbitrary primers targeting the chromosome. PCR products were sequenced, and insertion sites were determined by sequence homology searches against the *A. hydrophila* genome sequence.

### Quantification of colony opacity

To evaluate colony opacity, overnight cultures were spotted onto LB agar plates and incubated at 37°C for 12 h. Colony images were captured under identical conditions, and colony opacity/translucency was quantified using ImageJ software as described previously [25, 26]. Relative values were calculated by setting the WT value to 100. The assay was performed independently three times.

Analysis of secreted proteins by SDS-PAGE, silver staining, and Western blotting

WT, OS8, and OS9 were cultured in 5 mL of LB broth at 37°C with shaking overnight. Culture turbidity was measured at OD₆₀₀, and each culture was adjusted to the same optical density. The normalized cultures were inoculated into fresh LB broth and incubated for an additional 3 h at 37°C with shaking. After incubation, cultures were readjusted to OD₆₀₀ = 1.0 and centrifuged at 10,000 rpm for 10 min at room temperature to separate culture supernatants and bacterial pellets. For preparation of secreted protein samples, culture supernatants were precipitated with trichloroacetic acid (TCA) on ice for 20 min and centrifuged at 12,000 rpm for 20 min at 4°C. The resulting pellets were washed with cold diethyl ether, centrifuged again at 12,000 rpm for 20 min at 4°C, air-dried, and resuspended in SDS sample buffer. Bacterial pellets were directly resuspended in SDS sample buffer. Samples were separated by SDS-PAGE using a molecular weight marker (FastGene™ BlueStar Prestained Protein Marker, Nippon Genetics, NE-MWP03) and visualized by silver staining using the Silver Stain II kit (FUJIFILM Wako Pure Chemical Corporation, Osaka, Japan).

For detection of FLAG-tagged AerA, prepared samples were separated by SDS-PAGE, transferred onto membranes, and probed with monoclonal ANTI-FLAG M2 antibody (Sigma-Aldrich, St. Louis, MO, USA).

### Hemolysis assay

For qualitative evaluation of hemolysis, bacterial strains were cultured in 5 mL of LB broth at 37°C with shaking for 15 h. Cultures were adjusted to OD₆₀₀ = 1.0, serially diluted 10-fold, and 1.0 µL of each dilution was spotted onto sheep blood agar plates. After incubation at 37°C for at least 12 h, hemolytic zones were examined.

For quantitative hemolysis assays, bacterial cultures were prepared under the same conditions and separated into culture supernatants and bacterial suspensions. Mouse erythrocytes were prepared as a 2% suspension in phosphate-buffered saline (PBS) after careful removal of plasma and buffy coat. Aliquots (50 µL) of the erythrocyte suspension were dispensed into 96-well plates and mixed with 50 µL of either culture supernatant or bacterial suspension. After incubation at 37°C for 30 min, plates were centrifuged at 2,400 rpm for 10 min, and the supernatants were transferred to new plates. Absorbance was measured at 540 nm. Complete hemolysis was defined using distilled water, and nonhemolytic controls were prepared using PBS. Hemolytic activity was expressed as a percentage relative to the complete hemolysis control.

### Animals

Five-week-old female C57BL/6 and BALB/c mice were purchased from Japan Jackson Laboratory and housed under specific-pathogen-free conditions with free access to food and water on a 12-h light/dark cycle. C57BL/6 mice were used for all animal experiments except in vivo bioluminescence imaging, for which BALB/c mice were used. All animal experiments were approved by the Institutional Animal Care and Use Committee of Kitasato University and were performed in accordance with the guidelines of the Japanese Association for Laboratory Animal Science (JALAS) (approval no. 24-033). This study is reported in accordance with the ARRIVE guidelines.

### Mouse NSTI model

Bacteria cultured at 37°C were diluted 1:100 in fresh medium and incubated for an additional 8 h. Cultures were collected by centrifugation at 7,000 × *g* for 3 min, washed once with PBS, and resuspended in the same medium. Each mouse was inoculated subcutaneously in the right thigh with 1 × 10⁷ CFU in 100 µL (n = 6 per group unless otherwise indicated). Mice were anesthetized with isoflurane (DS Pharma Animal Health, Japan) during infection procedures and monitored hourly. Animals that reached predefined humane endpoints were euthanized by sevoflurane overdose (Nikko Pharmaceutical Co., Ltd., Japan) under deep anesthesia, in accordance with AVMA guidelines.

### Serum biomarker measurements

At 10 h post-infection, mice were deeply anesthetized and euthanized, and whole-blood samples were collected by cardiac puncture. Blood samples were centrifuged at 1,200 × *g* for 30 min to obtain serum. Serum lactate dehydrogenase (LDH), creatine phosphokinase (CPK), and aspartate aminotransferase (AST) concentrations were measured using a Dimension EXL with the LM Integrated Chemistry System (Siemens, Tokyo, Japan).

### Histological examination

At 10 h post-infection, infected thighs were harvested, decalcified, and embedded in paraffin. Sections (4 µm thick) containing both subcutaneous and muscle tissues were prepared and stained with hematoxylin and eosin (H&E). The stained sections were examined and photographed using a light microscope.

### Enumeration of bacterial burdens in muscle and in spleen

At 12 h after subcutaneous infection with *A. hydrophila*, infected thigh tissues and spleens were harvested. Muscle tissue directly beneath the inoculation site was dissected, homogenized in 1 mL of 0.1% gelatin in PBS for 5 s using an IKA EUROSTAR homogenizer (IKA Werke, Germany) at 1,300 rpm, and centrifuged at 800 rpm for 5 min. Supernatants were serially diluted 10-fold and plated in duplicate on LB agar plates. After incubation at 37°C for 24 h, colony-forming units (CFU) were enumerated. Bacterial burdens in spleens were determined in the same manner.

### In vivo bioluminescence imaging

For in vivo bioluminescence imaging, BALB/c mice were used. Bioluminescent WT and mutant strains were visualized using an IVIS-200 imaging system (PerkinElmer, Waltham, MA, USA). Mice were anesthetized with isoflurane, and images were acquired with a fixed exposure time of 1 min at 3, 6, 9, and 12 h after subcutaneous infection. The imaging angle was modified from our previous *V. vulnificus* protocol to better capture bacterial spread within soft tissues [27]. Imaging experiments were performed using three independent mice per strain.

### Statistical analysis

All statistical analyses were performed using GraphPad Prism version 8 (GraphPad Software, San Diego, CA, USA). Comparisons between two groups were analyzed using the Mann-Whitney *U* test. Comparisons among more than two groups were analyzed using the Holm-Sidak multiple-comparisons test, with each group compared with WT. Survival curves were compared using the log-rank (Mantel-Cox) test. Data are presented as means ± standard error of the mean (SEM). A *P* value of <0.05 was considered statistically significant.

## RESULTS

### T2SS-related mutants show reduced colony opacity and attenuated necrosis

During phenotypic screening of an *A. hydrophila* transposon insertion mutant library, we identified mutants exhibiting altered colony opacity compared with the wild-type (WT) strain. Bacterial suspensions were spotted onto LB agar plates, and colonies formed after 12 h of incubation were evaluated based on their opacity, with translucent variants selected for further analysis. While WT colonies appeared opaque, the Tn mutants OS3, OS4, OS7, OS8, OS9, OS14, and OS15 exhibited reduced colony opacity (Fig. 1A). To identify the disrupted genes in these mutants, transposon insertion sites were determined (Table 1). This analysis revealed that OS8 harbored a transposon insertion in *gspD*, whereas OS9 carried an insertion in *gspC*. Accordingly, OS8 and OS9 were designated the *gspD* mutant and the *gspC* mutant, respectively, in the following analyses (Fig. 1B). In a mouse NSTI model, WT-infected mice developed severe soft tissue necrosis at the infection site, whereas OS8-infected mice showed only mild necrosis and OS9-infected mice showed little to no necrosis (Fig. 1C). These results indicate that the GspCD-dependent T2SS contributes to *A. hydrophila* soft tissue pathogenesis.

**Figure 1.**
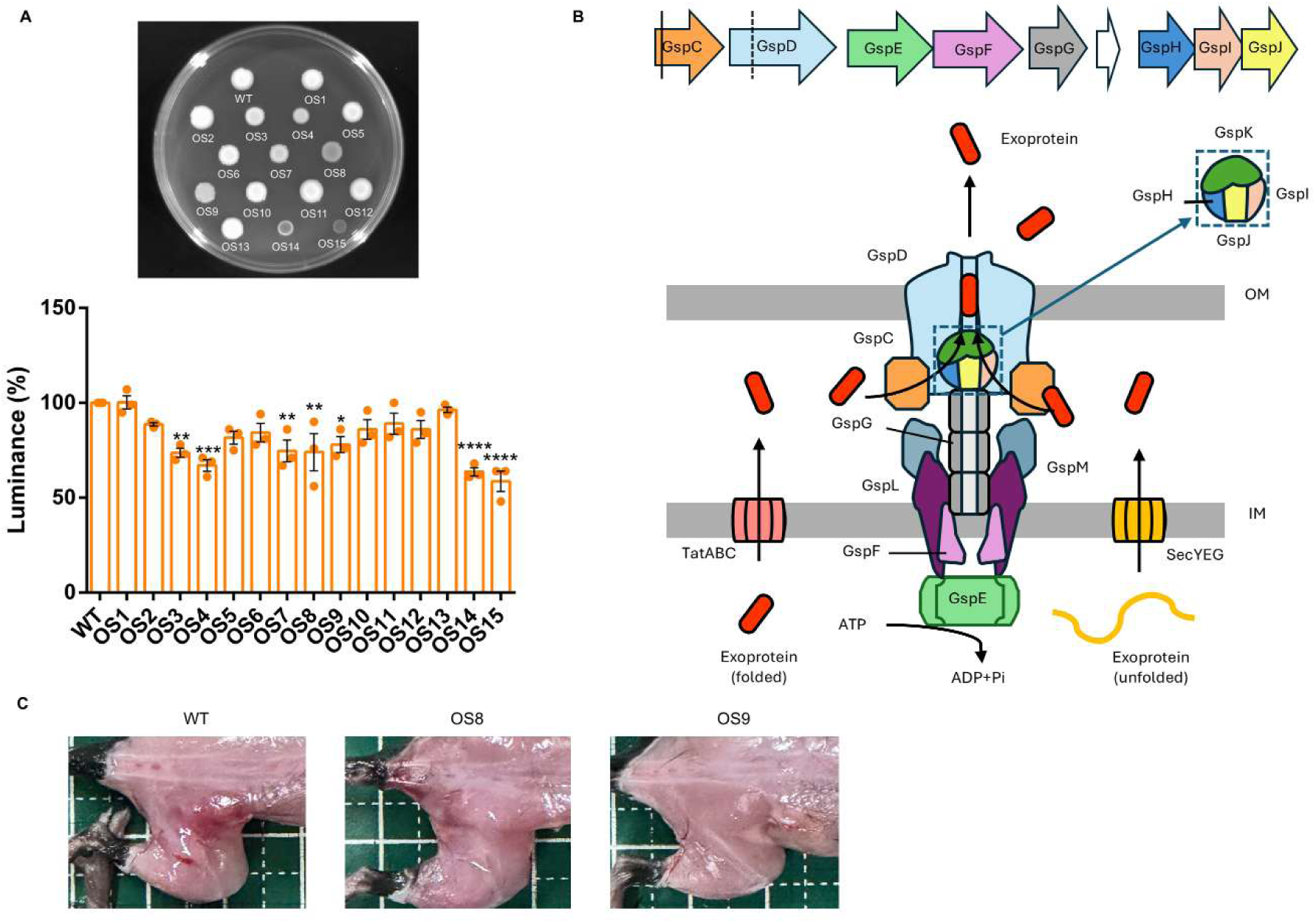
Phenotypic screening of transposon mutants (A) Representative images of colony morphology of the wild-type (WT) strain and transposon mutants (OS1-OS15) grown on LB agar plates. Quantification of colony opacity is shown as relative values normalized to WT (%). Data represent mean ± SEM (n = 3). The image is representative of three independent experiments. Statistical analysis was performed using the Holm-Sidak multiple-comparisons test, with each group compared with WT. \**P* < 0.05; \*\**P* < 0.01; \*\*\**P* < 0.001; \*\*\*\**P* < 0.0001. (B) Schematic representation of transposon insertion sites in selected mutants and the organization of the type II secretion system (T2SS). Insertions in *gspC* and *gspD* are indicated. (C) In vivo screening in a murine NSTI model. Representative gross images of infected tissues following infection with WT, OS8, or OS9.

**Table 1.**
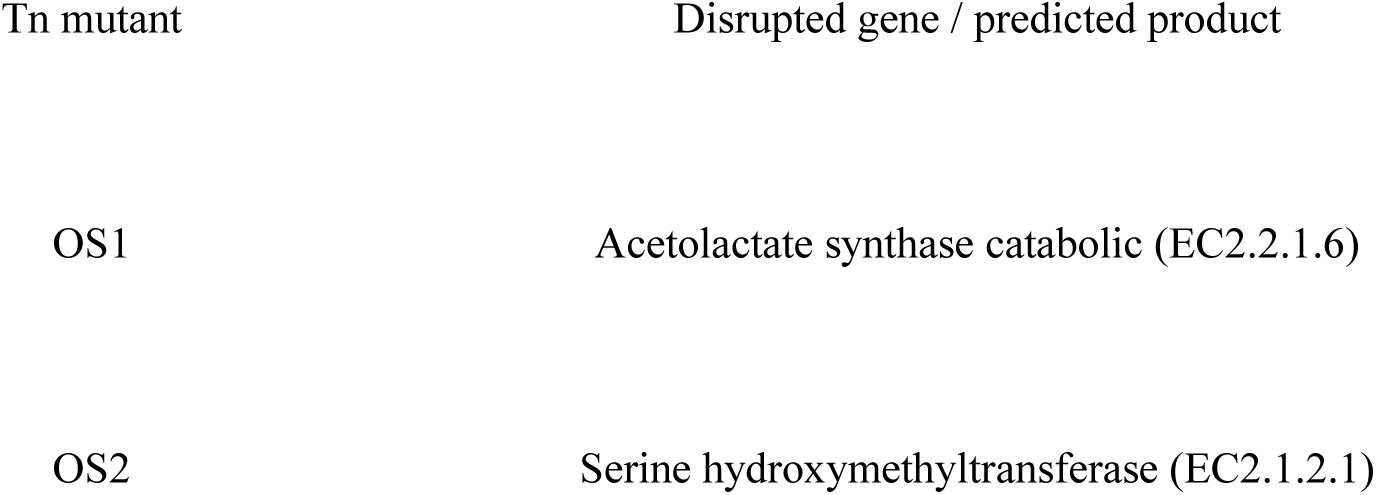
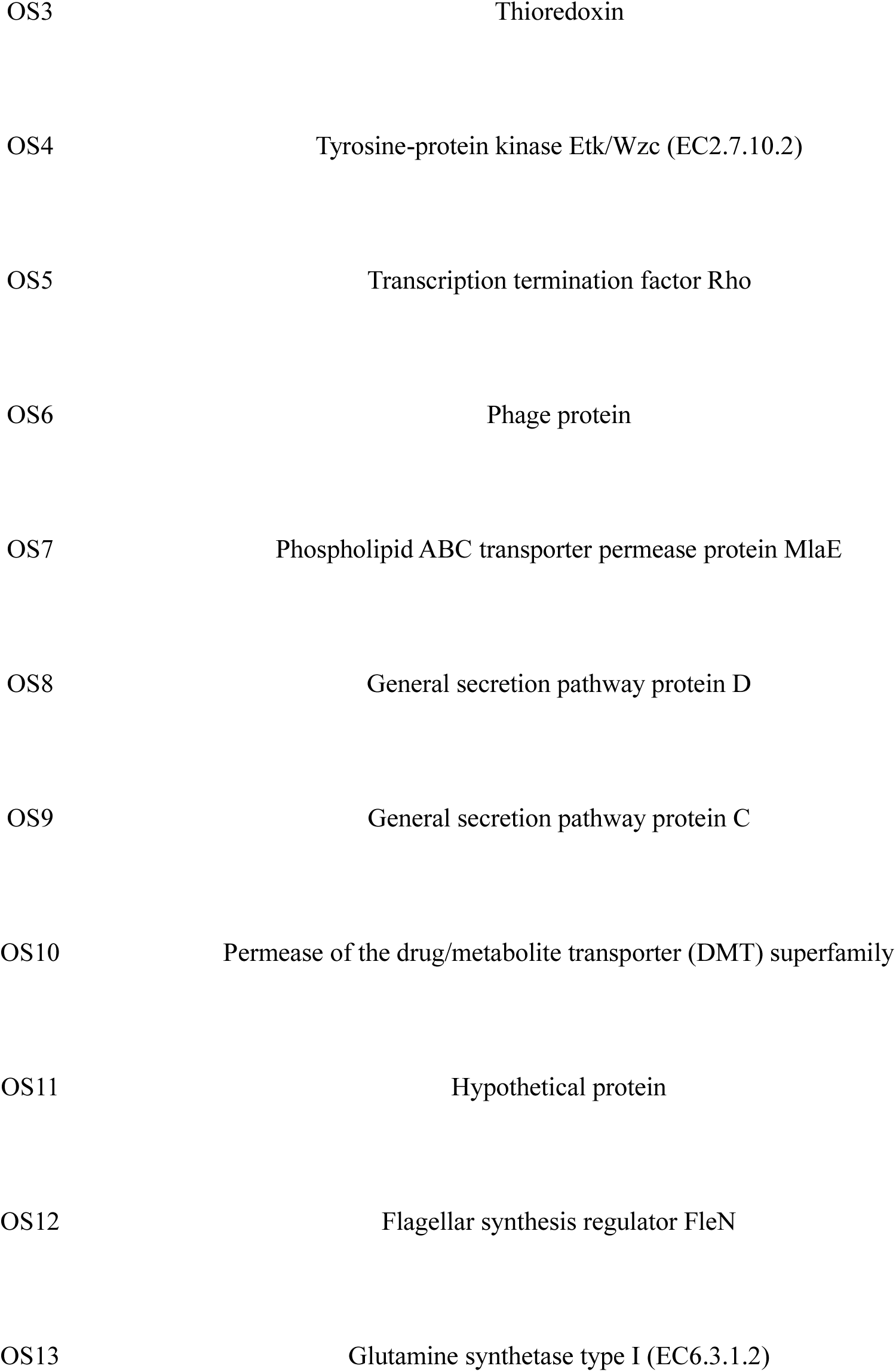
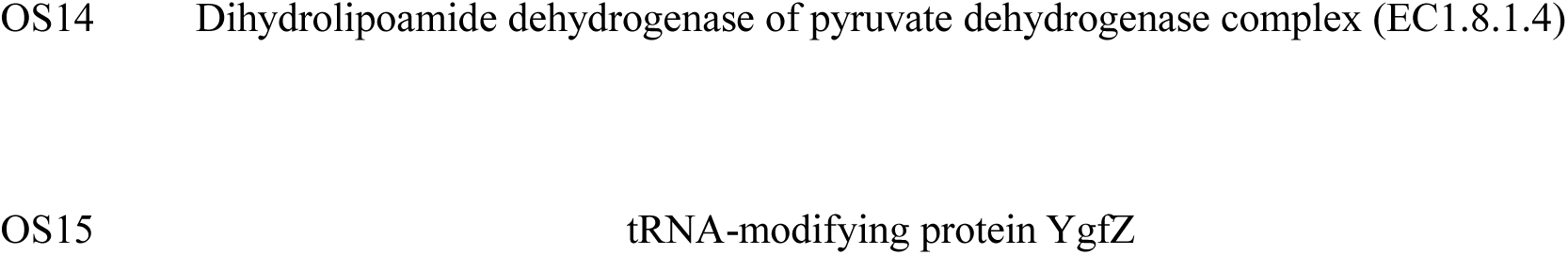
Genes disrupted by transposon insertion in screened mutants.

### Disruption of *gspC* alters extracellular protein secretion and composition

Given that GspC and GspD are essential components of the type II secretion system (T2SS), we hypothesized that T2SS-dependent secretion would be impaired in OS8 and OS9. To assess T2SS function, a FLAG-tagged truncated AerA construct corresponding to the secretion signal and N-terminal region of AerA (predicted molecular mass, approximately 20 kDa) was expressed. FLAG-reactive signals were detected in the culture supernatant of WT, whereas they were absent from the supernatants of OS8 and OS9 (Fig. 2A), indicating defective secretion of the truncated AerA construct in these mutants.

**Figure 2.**
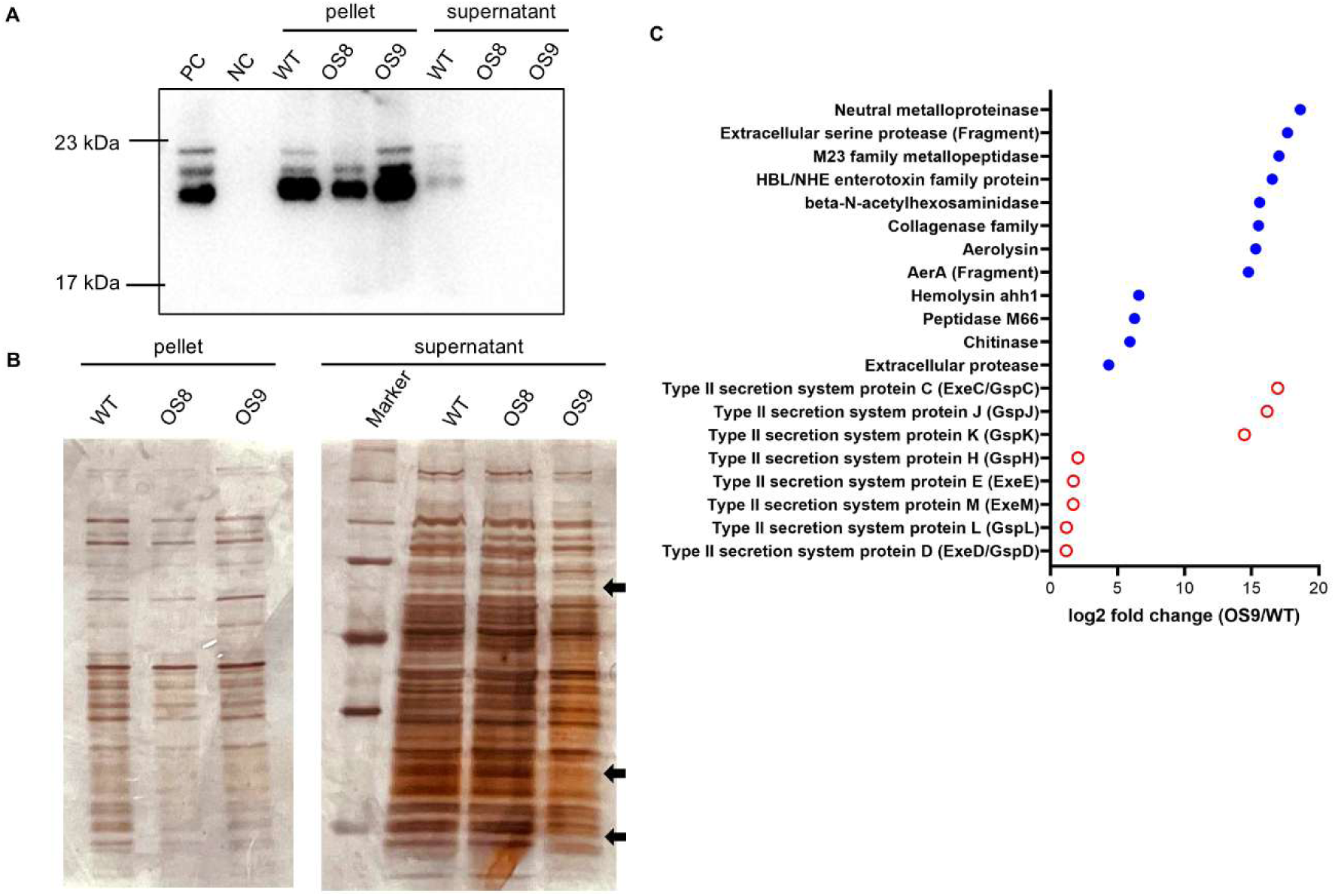
Analysis of extracellular protein secretion (A) Western blot analysis of FLAG-tagged AerA in culture supernatant and pellet fractions from WT, OS8, and OS9. PC, E. coli DH5α carrying the Aerolysin expression plasmid; NC, WT *A. hydrophila* without the plasmid. (B) Silver staining of proteins in culture supernatant and pellet fractions from WT, OS8, and OS9. (C) Representative proteins with reduced abundance in the culture supernatant of the *gspC* mutant (OS9) relative to WT. Proteins are plotted by log2 fold change (OS9/WT). Filled blue circles indicate representative virulence-associated extracellular proteins, and open red circles indicate T2SS-related proteins.

To further examine extracellular protein profiles, bacterial cultures were separated into supernatant and pellet fractions and analyzed by silver staining. No notable differences were observed between WT and OS8 in either fraction, whereas three bands present in the WT and OS8 supernatants were absent from the OS9 supernatant (Fig. 2B), suggesting a greater alteration in extracellular protein composition in the gspC mutant.

Proteomic analysis of culture supernatants from WT and OS9 was then performed to further define these differences. Although the total protein amounts were comparable (Sup Fig. 1), 119 proteins were reduced in OS9 compared with WT (Table S1). Representative proteins reduced in OS9 included several known or predicted T2SS-associated extracellular factors, such as AerA, Ahh, lipase, metalloprotease, collagenase-related proteins, and extracellular serine protease, as well as multiple T2SS-related proteins (Fig. 2C). Together, these results indicate that disruption of *gspC* broadly alters extracellular protein secretion and composition in *A. hydrophila*.

### T2SS-dependent secretion contributes to hemolytic activity

We next asked whether disruption of the T2SS affects hemolytic activity. Hemolytic zones were first assessed on blood agar plates. WT formed clear hemolytic zones, whereas no hemolytic zones were observed for OS8 or OS9 (Fig. 3A). Hemolytic activity was restored in the complemented strains of both mutants.

**Figure 3.**
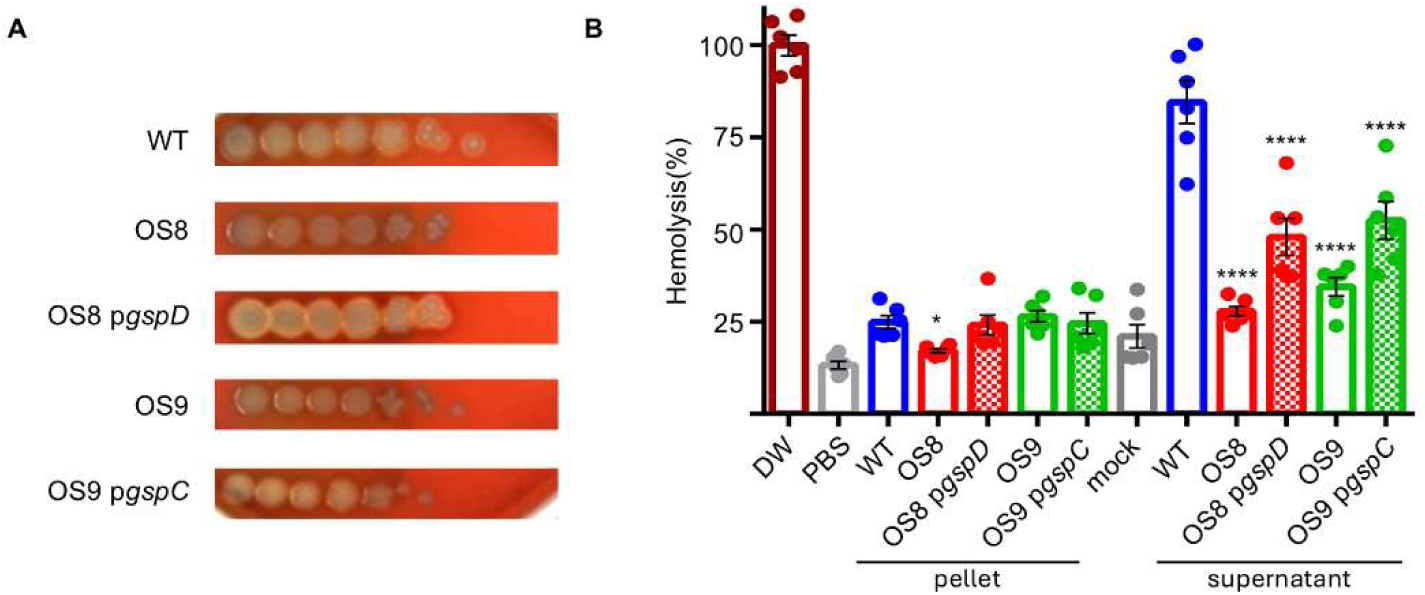
Hemolytic activity of T2SS mutants (A) Hemolysis on blood agar plates by the WT, OS8, OS9 and complemented strains. Representative images from three independent experiments are shown. (B) Quantification of hemolytic activity using mouse erythrocytes. Culture supernatant and pellet fractions from each strain were incubated with erythrocytes, and hemolysis was measured by absorbance at 540 nm. Data represent mean ± SEM (n = 6). Statistical analysis was performed using the Holm-Sidak multiple-comparisons test, with each group compared with WT. \**P* < 0.05; \*\*\*\**P* < 0.0001.

We further quantified hemolytic activity using mouse erythrocytes. No significant differences were observed among strains in the pellet fraction; however, hemolytic activity in the culture supernatants of OS8 and OS9 was significantly reduced compared with WT, and this reduction was restored in the complemented strains (Fig. 3B). These findings indicate that T2SS-dependent extracellular factors contribute to hemolytic activity in *A. hydrophila* and support the conclusion that disruption of *gspD* or *gspC* causes the hemolysis-defective phenotype.

### T2SS-dependent secretion contributes to soft tissue damage in vivo

We next examined the contribution of T2SS-dependent secretion to tissue damage in the NSTI model. Serum levels of CPK, LDH, and AST/GOT were measured in infected mice. All three markers were elevated in WT-infected mice compared with non-infected controls, whereas mice infected with OS8 or OS9 showed lower values than WT-infected mice (Fig. 4A).

**Figure 4.**
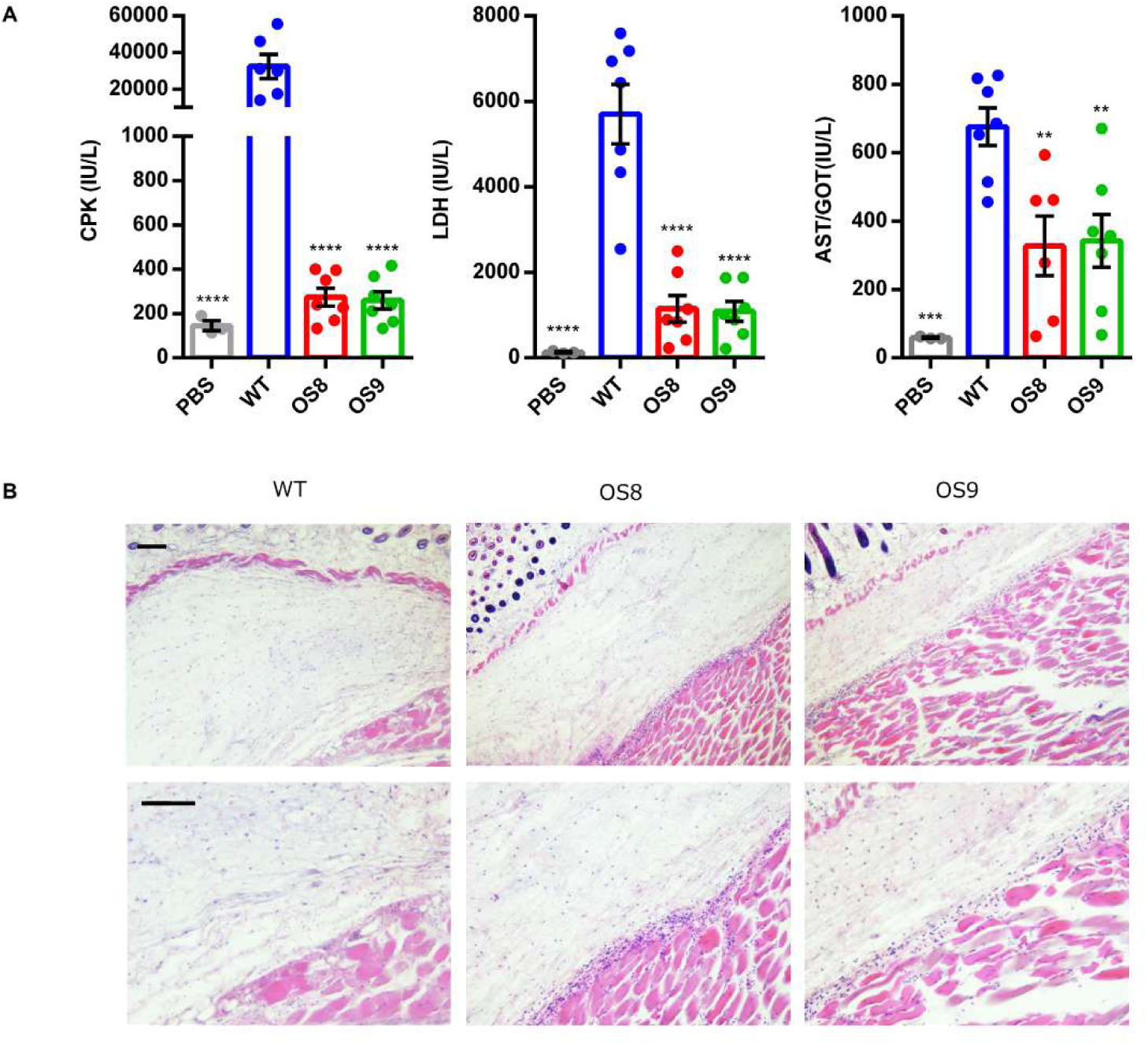
Soft tissue damage in a murine NSTI model (A) Serum levels of CPK, LDH, and AST in mice infected with WT, OS8, or OS9. Data represent mean ± SEM (n = 6). Statistical analysis was performed using the Holm-Sidak multiple-comparisons test, with each group compared with WT. \*\**P* < 0.01; \*\*\**P* < 0.001; \*\*\*\**P* < 0.0001. (B) Histopathological analysis of infection sites (thigh muscle) from mice infected with WT, OS8, or OS9. Representative H&E-stained sections are shown. The upper panels show low-magnification views, and the lower panels show higher-magnification views of the same sections. All scale bars (100 µm) are shown in the leftmost panels.

Histopathological analysis of infected tissues at 10 h post-infection revealed marked neutrophil infiltration and degeneration and destruction of muscle cells in WT-infected mice (Fig. 4B). In contrast, OS8-infected mice showed only mild neutrophil infiltration and cellular damage, whereas OS9-infected mice exhibited minimal pathological changes (Fig. 4B). These histopathological findings were consistent with the gross pathological changes observed at the infection site (Fig. 1C). Collectively, these results indicate that T2SS-dependent extracellular factors contribute substantially to soft tissue damage during *A. hydrophila* infection.

### T2SS-dependent secretion promotes in vivo expansion and dissemination

To evaluate the impact of T2SS-dependent tissue damage on bacterial behavior in vivo, bioluminescent *A. hydrophila* strains carrying chromosomally integrated *luxCDABE* were used to monitor bacterial expansion and dissemination by in vivo imaging system (IVIS) analysis. In WT-infected mice, signal intensity increased after 6 h post-infection and spread beyond the inoculation site. In contrast, signal intensity decreased after 6 h in OS8- and OS9-infected mice, with signals nearly disappearing by 12 h post-infection (Fig. 5A).

**Figure 5.**
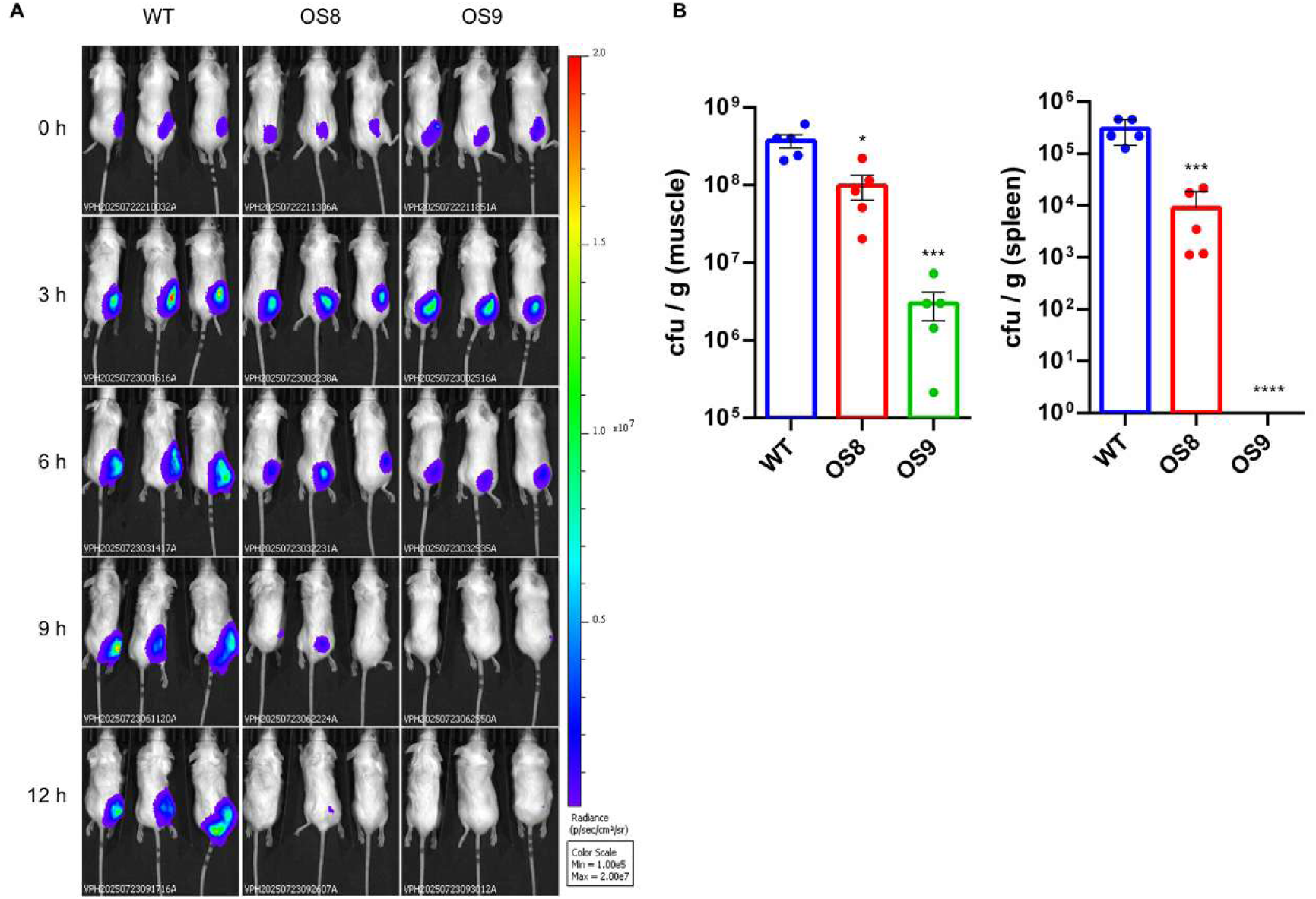
In vivo bacterial proliferation and dissemination (A) Bioluminescent signals of WT, OS8, and OS9 were monitored at the indicated time points following subcutaneous infection in a murine NSTI model. Images were acquired using identical imaging parameters. Representative bioluminescence images are shown (n = 3 per group). (B) Bacterial burdens in muscle at the infection site and in the spleen of mice infected with WT, OS8, or OS9. Data represent mean ± SEM (n = 6). Statistical analysis was performed using the Holm-Sidak multiple-comparisons test, with each group compared with WT. \**P* < 0.05; \*\*\**P* < 0.001; \*\*\*\**P* < 0.0001.

To assess local expansion and systemic dissemination, bacterial burdens were quantified in muscle tissue and in the spleen. WT-infected mice showed higher bacterial burdens in both tissues than mice infected with OS8 or OS9 (Fig. 5B). Moreover, OS9 was more strongly attenuated than OS8, and no bacteria were detected in the spleens of OS9-infected mice (Fig. 5B). These results indicate that T2SS-dependent secretion promotes bacterial expansion at the infection site and dissemination to systemic organs in vivo.

### T2SS-dependent secretion is critical for host lethality

Finally, we assessed the contribution of T2SS-dependent secretion to host lethality using the mouse NSTI model. All mice infected with WT died within 20 h post-infection (Fig. 6). In contrast, some mice infected with OS8 survived throughout the observation period, whereas OS9 caused no deaths (Fig. 6). These results show that T2SS-dependent secretion is critical for the lethal outcome of *A. hydrophila* soft tissue infection.

**Figure 6.**
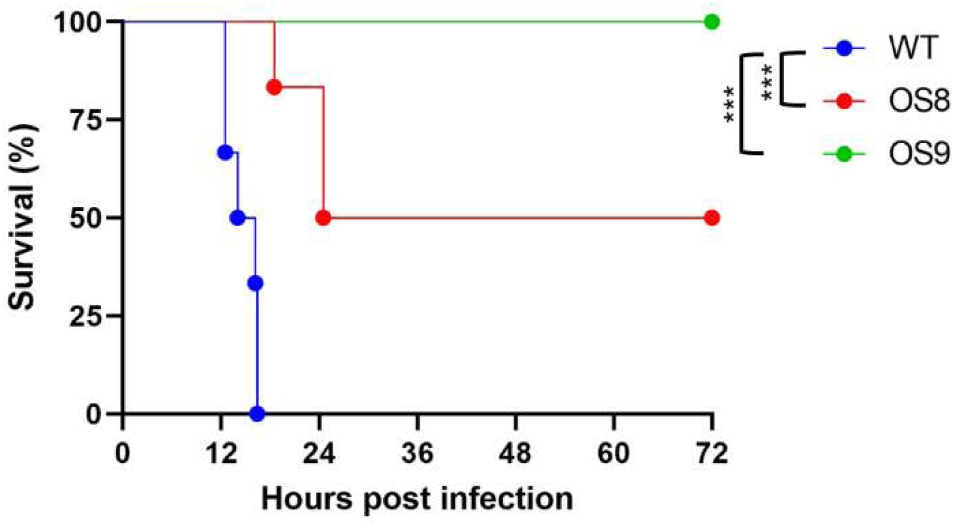
Survival of mice infected with T2SS mutants Survival curves of mice infected with WT, OS8, or OS9. Mice were monitored over the indicated period following subcutaneous infection in a murine NSTI model (n = 6 per group). Statistical analysis was performed using the log-rank (Mantel–Cox) test. \*\*\**P* < 0.001.

## DISCUSSION

*Aeromonas hydrophila* causes necrotizing soft tissue infections (NSTIs), a severe disease characterized by rapid progression and high mortality [4, 8]. In this study, we identify the GspCD-dependent type II secretion system (T2SS) as a central determinant of *A. hydrophila* pathogenesis in NSTIs. Disruption of *gspC* or *gspD* impaired secretion of a FLAG-tagged truncated AerA construct, reduced hemolytic activity, attenuated tissue damage, decreased in vivo expansion and dissemination, and markedly reduced lethality. These findings indicate that the T2SS contributes not only to extracellular protein secretion but also to the progression of invasive and lethal soft tissue infection.

The initial screen was based on colony opacity, which likely reflects changes in extracellular structures or secreted products (Fig. 1A). Using this phenotype, we identified mutants with transposon insertions in *gspD* and *gspC*. Both OS8 and OS9 were impaired in T2SS-dependent secretion, as demonstrated by the loss of FLAG-tagged AerA signals in culture supernatants (Fig. 2A). Because Aerolysin and hemolysins are membrane-active pore-forming toxins [5, 12, 13, 28], the reduced hemolytic activity observed in both mutants is consistent with impaired secretion of these factors. Restoration of hemolysis by complementation further supports the requirement of *gspD* and *gspC* for this phenotype (Fig. 3A and 3B). Importantly, proteomic analysis extended this model beyond a single secreted factor by showing that multiple known or predicted extracellular virulence-associated proteins were reduced in the supernatant of the *gspC* mutant, including AerA, Ahh, lipase, metalloprotease, collagenase-related proteins, and extracellular serine protease, together with several T2SS-related proteins (Fig. 2C and Table S1). These findings indicate that disruption of GspC broadly alters the extracellular protein landscape associated with T2SS-dependent secretion.

The in vivo data directly link these secretion defects to disease pathogenesis [14, 24]. In the NSTI model, WT infection caused marked increases in serum CPK, LDH, and AST, together with pronounced neutrophil infiltration and severe muscle damage (Fig. 4A and 4B). These changes were attenuated in the *gspD* mutant and were minimal in the *gspC* mutant, consistent with the gross pathological changes observed at the infection site. In addition, WT showed robust expansion at the infection site and dissemination to the spleen, whereas both mutants were markedly impaired in these processes, and no bacteria were detected in the spleens of mice infected with OS9 (Fig. 5B). Together with the IVIS results, these findings indicate that T2SS-dependent secretion contributes not only to host tissue injury but also to bacterial expansion and dissemination in vivo. We previously showed that motility is required for efficient spread of *A. hydrophila* in soft tissues [16]. The present study extends this model by demonstrating that, in addition to motility, T2SS-dependent extracellular factors are required for efficient tissue destruction, local expansion, systemic spread, and lethal outcome during NSTIs. Thus, NSTI progression appears to depend on coordinated interactions between bacterial motility and secretion-mediated remodeling of the infected tissue environment.

Notably, the *gspC* mutant showed stronger attenuation than the *gspD* mutant across multiple assays. In many Gram-negative bacteria, GspD is considered the key structural component of the T2SS because it forms the outer membrane secretin channel [29–34], whereas GspC is generally thought to function in substrate recognition and linkage of the secretion machinery [14, 20, 32, 35]. Our findings suggest that GspC may have a particularly important role in shaping the extracellular output associated with T2SS-dependent virulence in *A. hydrophila*. Although the mechanistic basis for this difference remains to be determined, the stronger phenotype of the *gspC* mutant raises the possibility that disruption of GspC causes broader functional consequences for secretion-associated processes than disruption of GspD under the conditions tested here.

In summary, our findings establish the GspCD-dependent T2SS as a central contributor to *A. hydrophila* NSTI pathogenesis. Rather than acting through a single toxin alone, this secretion system coordinates the extracellular export of multiple proteins associated with hemolysis, tissue destruction, in vivo expansion, dissemination, and host mortality. These results highlight the importance of T2SS-dependent extracellular protein organization in severe soft tissue infection and provide a framework for understanding how secretion-system-level perturbations reshape bacterial virulence during invasive disease.

## DATA AVAILABILITY STATEMENT

The datasets generated and analyzed during the current study, including raw western blot images, raw numerical data, and supplementary tables, are openly available in Zenodo at https://doi.org/10.5281/zenodo.19400541 [36].

## ETHICS APPROVAL

All animal experiments were approved by the Institutional Animal Care and Use Committee of Kitasato University and were performed in accordance with the guidelines of the Japanese Association for Laboratory Animal Science (JALAS) (approval no. 24-033). This study is reported in accordance with the ARRIVE guidelines.

## FUNDING

This work was supported by Grants-in-Aid for Scientific Research (KAKENHI) from the Japan Society for the Promotion of Science (JSPS), Grant Numbers 22K14998 and 22KK0289 awarded to Kohei Yamazaki.

## CONFLICTS OF INTEREST

No potential conflict of interest was reported by the author(s).

## ACKNOWLEDGMENTS

The authors thank ChatGPT (OpenAI) and the Nature Research Editing Service for assistance with English language editing and manuscript polishing. These tools were used solely to improve clarity and readability of the text and did not influence the study design, data analysis, or interpretation of results.

